# One-step generation of mice by Transposon-Enhanced Multi-Plex Orchestration Editing(TEMPO-Editing)

**DOI:** 10.64898/2026.07.29.741479

**Authors:** Maiko Inotsume, Kasumi Yumoto, Tomoki Chiba, Masayasu Sega, Takahide Matsushima, Hiroshi Asahara

## Abstract

Genome-edited mice are widely used to elucidate molecular mechanisms in vivo and are an indispensable tool, particularly for studies aimed at clarifying gene function at the organismal level. Currently, there is an increasing demand for mice in which multiple genes are simultaneously modified in order to investigate interactions among multiple genes. Notably, generating conditional multiple-gene knockout mice with temporal and spatial specificity requires extensive crossing between multiple Cre-driver mice and floxed mice, resulting in a prolonged time frame for line establishment.

To address this limitation, we developed Transposon-Enhanced Multi-Plex Orchestration Editing (TEMPO-editing), a single-step strategy for generating multiple-gene-edited mice. TEMPO-editing enables simultaneous modification of multiple genes through the integration of transposon, Cre-loxP, and CRISPR/Cas9 systems. Using the DNA transposon piggyBac, we constructed a single cassette harboring gRNAs targeting genes of interest together with a conditionally expressed Cas9 (lsl-Cas9). By injecting this cassette into fertilized eggs of Cre mice, we enabled the generation of temporally and spatially specific genome edited mice in the F0 generation.

In this study, we generated double-gene-edited mice targeting *Hoxa13* and *Hoxd13*, which are key regulators of embryonic body patterning and are essential for autopod development. The phenotype observed in these mice was consistent with the incomplete autopod phenotype previously reported in mice generated by crossing *Hoxa13* knockout and *Hoxd13* knockout mice. These results demonstrate the utility of TEMPO-editing for the generation of multiple-gene-edited mice in the F0 generation. Notably, this study establishes a simplified strategy for producing conditional multi-gene-edited mice, a process that has traditionally required substantial time and labor using conventional methods.

## Introduction

Genetically engineered mice have become indispensable tools in modern molecular biology and life science research. In particular, they are widely used to investigate gene functions essential for maintaining biological homeostasis by disrupting genes critical for survival, as well as to model pathological conditions through the modification of disease-associated genes, thereby facilitating studies aimed at elucidating disease mechanisms and developing preventive and therapeutic strategies. However, the generation of genetically modified mouse lines requires substantial time, labor, and financial investment, making the establishment of even a single mouse strain a considerable burden for research.

Traditionally, genetically modified mice have been generated using embryonic stem (ES) cell–based technologies, which were established following the derivation of ES cells in 1981 and subsequently adapted for genetic manipulation [1–4]. In this approach, genetically modified ES cells are injected into mouse blastocysts to generate chimeric mice, which are then bred over multiple generations to establish mouse lines carrying the desired genotype. Because of the requirement for germline transmission and successive breeding steps, this process typically takes several months to obtain the target mouse strain [5]. Subsequently, in 2012, the CRISPR/Cas9 system was developed by Charpentier and Doudna, marking a major breakthrough in genome editing technologies [6]. By simply introducing a guide RNA (gRNA) of approximately 20 nucleotides together with Cas9 mRNA or a Cas9 expression vector into fertilized eggs, targeted genetic modifications could be achieved within a single generation. In 2013, Huang et al. and Jaenisch et al. independently reported the successful generation of knockout mice using the CRISPR/Cas9 system [7–9]. However, despite these advances, the efficiency and reliability of genome editing using this approach were still limited, leaving room for further methodological improvements [10].

Thus, the generation of genome-edited mice remains challenging due to the extended time required for strain establishment and the need for advanced genome engineering techniques. To address these limitations, a variety of more efficient and reproducible genome editing strategies have been developed. For example, a method employing two guide RNAs together with a large DNA donor cassette flanked by loxP sites has been reported, enabling the simultaneous introduction of two loxP sequences in a single step [11]. More recently, Hatada and colleagues at Gunma University developed an innovative two-step electroporation strategy in which loxP sequences are introduced sequentially into each allele. In this approach, a single cleavage event occurs at the one-cell stage and another at the two-cell stage, thereby minimizing the risk of large deletions between the two gRNA target sites. In addition, electroporation represents a low-invasiveness delivery method, offering the advantage of reduced damage to embryos [12].

Nevertheless, even with these advanced techniques, the establishment of spatiotemporally controlled knockout mice targeting multiple genes often requires additional rounds of breeding, which may extend over several years. Therefore, from the perspective of accelerating and simplifying the generation of multi-gene–modified mouse models, substantial room remains for further methodological improvement and the development of novel genome editing strategies.

In recent years, novel genome editing technologies such as base editing and prime editing have also been developed [13,14]. However, these approaches are not universally applicable, as they are restricted in the types of genetic alterations that can be introduced and often suffer from limitations in editing efficiency and target scope. Therefore, from the perspectives of efficiency and versatility, genome editing strategies based on Cas9-mediated non-homologous end joining (NHEJ) continue to offer substantial potential for further development.

As proof of concept for the present study, we previously established a one-step method for generating conditional gene-edited mice by utilizing Tol2, a DNA transposon derived from medaka fish. In this approach, a gRNA expression cassette was introduced into fertilized eggs obtained from crosses between CAG-CreER and LSL-Cas9 mice, thereby markedly simplifying the multi-generational breeding processes required by conventional strategies [15].

Transposons were first described by Barbara McClintock in 1951 as genetic elements capable of relocating within the genome [16]. In current experimental animal research, DNA transposons such as Tol2 and piggyBac (PB), which excise DNA fragments flanked by terminal inverted repeats (ITRs) and integrate them into new genomic loci via a cut-and-paste mechanism, are widely used for genetic manipulation. Kawakami and colleagues were the first to report genome integration in vertebrates using a DNA transposon system based on Tol2 [17]. Building on this technology, Tol2- and piggyBac-based systems have since been extensively applied to genome editing and transgene delivery in mammalian models [18,19].

In this study, we refined our previously reported model [15] and focused on Hox genes, a group of transcription factors that play central roles in developmental processes. The Hox gene family consists of 39 members in the mouse genome and is critically involved in controlling pattern formation of the body axis and organ development during embryogenesis [20]. In particular, Hox genes belonging to the same paralogous group are known to function in a coordinated manner, acting cooperatively to regulate developmental programs [21]. In this study, we focused on *Hoxa13* and *Hoxd13*, two Hox genes that are essential for the formation of the distal limb (autopod), and attempted to generate double-gene-edited mice lacking both genes simultaneously. Previous studies have demonstrated that disruption of either *Hoxa13* or *Hoxd13* alone results in defective digit formation, whereas the combined loss of both genes leads to severe malformation of the entire autopod [22]. These double knockout mice were generated through ES cell–based targeted gene disruption followed by sequential breeding of individual knockout lines, a process that is presumed to require a substantial amount of time to establish the desired mouse strains. In this study, we induced region-specific deletion of *Hoxa13* and *Hoxd13* using *Prrx1*-Cre, a limb mesenchyme–specific Cre driver [23]. As a result, we successfully reproduced the autopod malformation phenotype previously reported for the combined loss of these genes. Taken together, this study presents a novel and efficient strategy for generating conditional multi-gene-edited mice, which substantially reduces the time and experimental burden traditionally associated with the production of such complex genetically engineered mouse models.

## Result

### TEMPO-Editing construction

We developed a genome-editing approach termed Transposon-Enhanced Multi-Plex Orchestration Editing (TEMPO-Editing). In this method, a vector containing multiple tandemly linked gRNAs, followed by a loxP-stop-loxP Cas9 (lsl-Cas9) cassette and mCherry, was introduced together with piggyBac transposase mRNA into fertilized embryos generated by in vitro fertilization (IVF) between male Cre-driver mice and wild-type females. The components were delivered by cytoplasmic microinjection into fertilized eggs. In this study, fertilized embryos were generated by in vitro fertilization (IVF) using male *Prrx1*-Cre mice, which express Cre recombinase specifically in limb mesenchymal cells. PiggyBac transposase mRNA was synthesized by in vitro transcription and co-injected into the cytoplasm of fertilized eggs together with a gRNA-expressing vector targeting *Hoxa13* and *Hoxd13* (*pCPX162_HoxAD13_pBSPB-CAG-LSL-Cas9-P2A-mCherry*). The following day, embryos that had developed to the two-cell stage were transferred into the oviducts of pseudopregnant recipient females, which subsequently delivered offspring approximately 20 days later (Fig. 1A).

**Figure 1.**
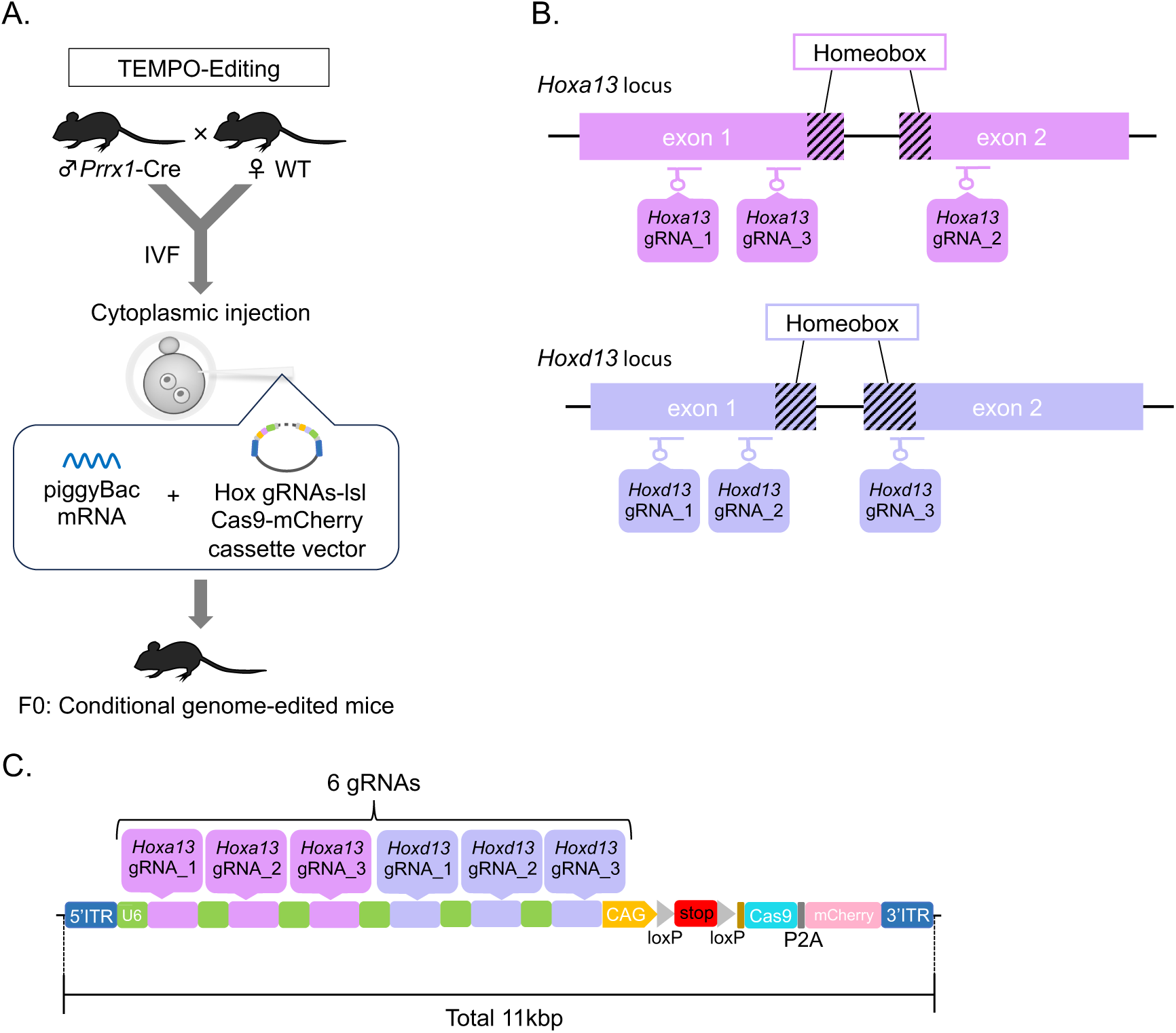
Generation of conditional *Hoxa13/Hoxd13* gene-edited mice using TEMPO-Editing. **(A)** Schematic overview of the TEMPO-Editing strategy. Male *Prrx1*-Cre mice were crossed with wild-type females, and fertilized embryos were obtained by in vitro fertilization (IVF). Fertilized embryos were subjected to cytoplasmic injection of piggyBac transposase mRNA together with a piggyBac donor vector carrying a Cre-dependent Cas9-mCherry cassette and multiplexed gRNAs targeting *Hoxa13* and *Hoxd13*. **(B)** Design of gRNAs targeting the *Hoxa13* and *Hoxd13* loci. Three gRNAs were designed for each gene to induce deletions within the highly conserved homeobox domain, thereby disrupting gene function. Colored boxes indicate exons, and hatched regions indicate the homeobox-coding region. **(C)** Structure of the piggyBac donor vector used for multiplex genome editing. The vector contains six U6 promoter-driven gRNA expression cassettes targeting *Hoxa13* (gRNA-1–3) and *Hoxd13* (gRNA-1–3). In addition, it includes a CAG promoter, a loxP-flanked transcriptional STOP cassette, and a Cas9-P2A-mCherry expression module. The total vector size is approximately 11 kb.

*Hoxa13* and *Hoxd13* play critical roles in limb development, particularly in the formation of the embryonic autopod. To target these genes, two target sites in exon 1 and one target site in exon 2 were designed for both *Hoxa13* and *Hoxd13*(Fig. 1B). A vector expressing three gRNAs targeting *Hoxa13* and three gRNAs targeting *Hoxd13* was constructed. Each gRNA expression unit consisted of a U6 promoter–gRNA–scaffold–terminator cassette, and six such units were tandemly assembled within a single vector. Downstream of the gRNA array, an lsl-Cas9 cassette was inserted, enabling Cre-dependent expression of Cas9 from the same genomic locus as the gRNAs. In addition, the fluorescent reporter mCherry was included to allow visualization of Cas9-expressing cells when necessary. To enable genomic integration via the piggyBac transposon system, the entire cassette containing the gRNAs, lsl-Cas9, and mCherry was flanked by piggyBac terminal repeat sequences. This design allows cut-and-paste integration of the transgene into the host genome through piggyBac-mediated transposition (Fig. 1C).

### *Hoxa13*/*Hoxd13* double-gene-edited mice

In mice carrying both the *Prrx1*-Cre transgene and the integrated gRNA cassette, incomplete formation of the autopod was observed, consistent with a previously reported phenotype [22]. Mice were photographed at birth (P0) or postnatal day 1.5 (P1.5) and again one week later (Fig. 2A). Genotyping was performed by PCR. Integration of the cassette was confirmed by amplification of the regions spanning 5′ ITR and sp2 and 3×FLAG and Cas9, respectively (Fig. 2B). Two independent microinjection experiments were performed. In the first experiment, 22 mice were born, of which 2 carried the integrated cassette, corresponding to an integration rate of 9.1%. Among these animals, only one mouse (Tg #1) also carried the Cre transgene, representing 4.5% of all offspring. In the second experiment, 4 mice were born, and one mouse (Tg #2) carried both the integrated cassette and the Cre transgene. Thus, 25% of the offspring carried both the cassette and Cre (Table 1).

**Figure 2.**
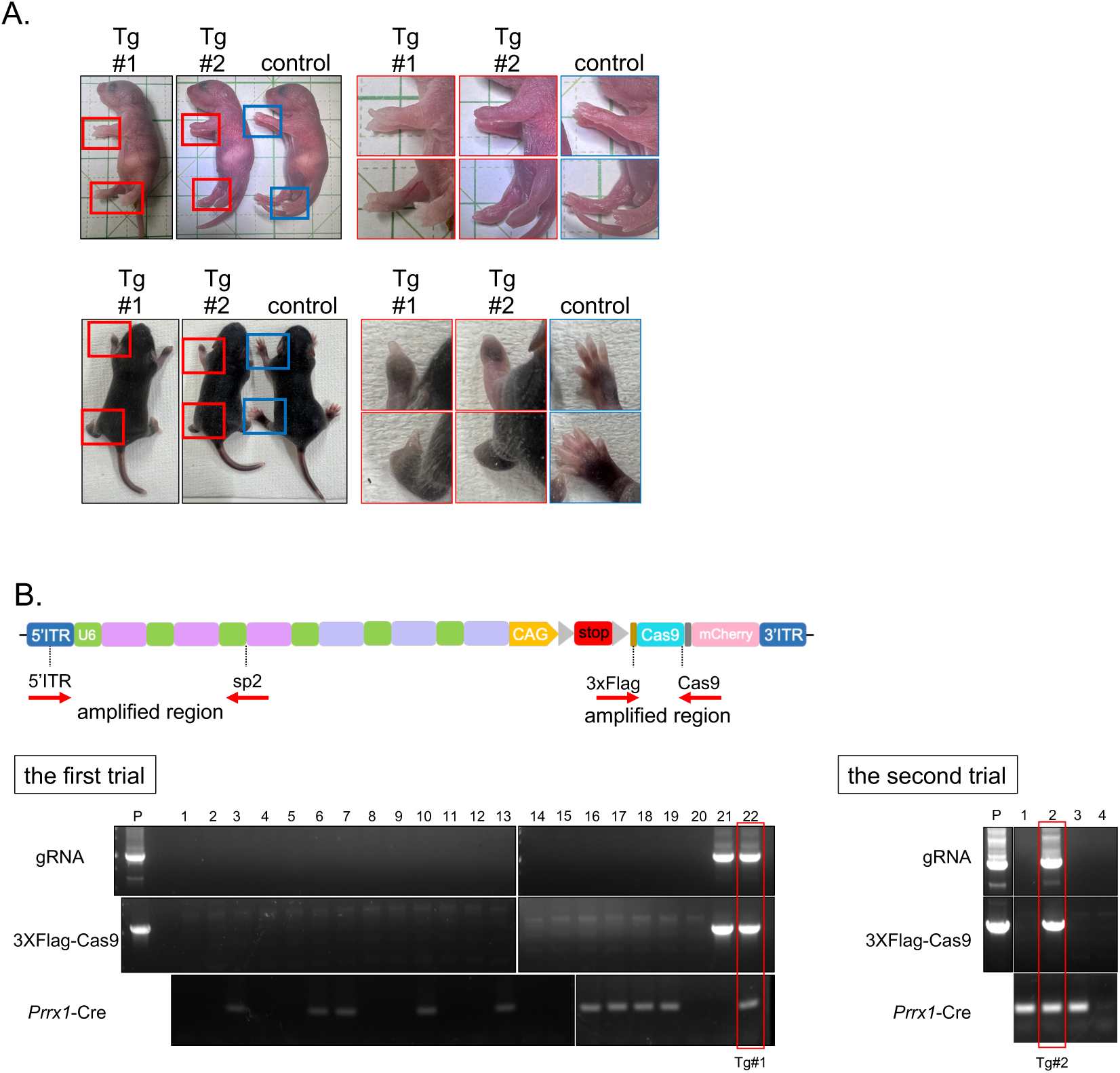
Phenotypic characterization and transgene validation of conditional *Hoxa13/Hoxd13* gene-edited mice generated by TEMPO-Editing. **(A)** Gross morphology of transgenic founder mice (Tg#1 and Tg#2) and control littermates. Whole-body images of neonatal mice and enlarged views of the forelimb and hindlimb autopods are shown. Both Tg#1 and Tg#2 exhibited morphological abnormalities in the autopod region compared with control mice. Red and blue boxes indicate the regions shown at higher magnification. **(B)** Genotyping analysis of transgenic founder mice. The upper panel shows the structure of the piggyBac donor vector and the positions of PCR primers used for transgene detection. Genomic DNA was analyzed by PCR to detect the gRNA cassette, the 3×Flag-Cas9 sequence, and the *Prrx1*-Cre transgene. P indicates the positive control (the donor vector used for embryo injection). Founder mice positive for the transgene are highlighted by red boxes and were identified as transgenic founder animals.

**Table 1.**

| No. of trial | No. of Microinjected Embryos | Transferred embryos | Births (% Transferred) | Transgenic pups (% Animals Born) [% Transferred] | Double-positive pups: Cre/gRNA cassettes (% Animals Born) [% Transferred] { % Transgenic born} |
| --- | --- | --- | --- | --- | --- |
| 1. | 299 | 99 | 22 (22.2%) | 2 (9.1%)[2.0%] | 1(4.5%)[1.0%]{50%} |
| 2. | 100 | 51 | 4(7.8%) | 1(25.0%)[2.0%] | 1(25.0%)[2.0%]{100%} |

### Verification of germline transmission of the gRNA cassette to the F1 generation

To evaluate germline transmission to the next generation, transgenic male mouse Tg #2 was used for in vitro fertilization (IVF) with wild-type (WT) female mice, and the transmission rate of the cassette to the offspring was assessed. Eight F1 mice were born, of which three displayed visually detectable incomplete autopod formation (Fig. 3A).

**Figure 3.**
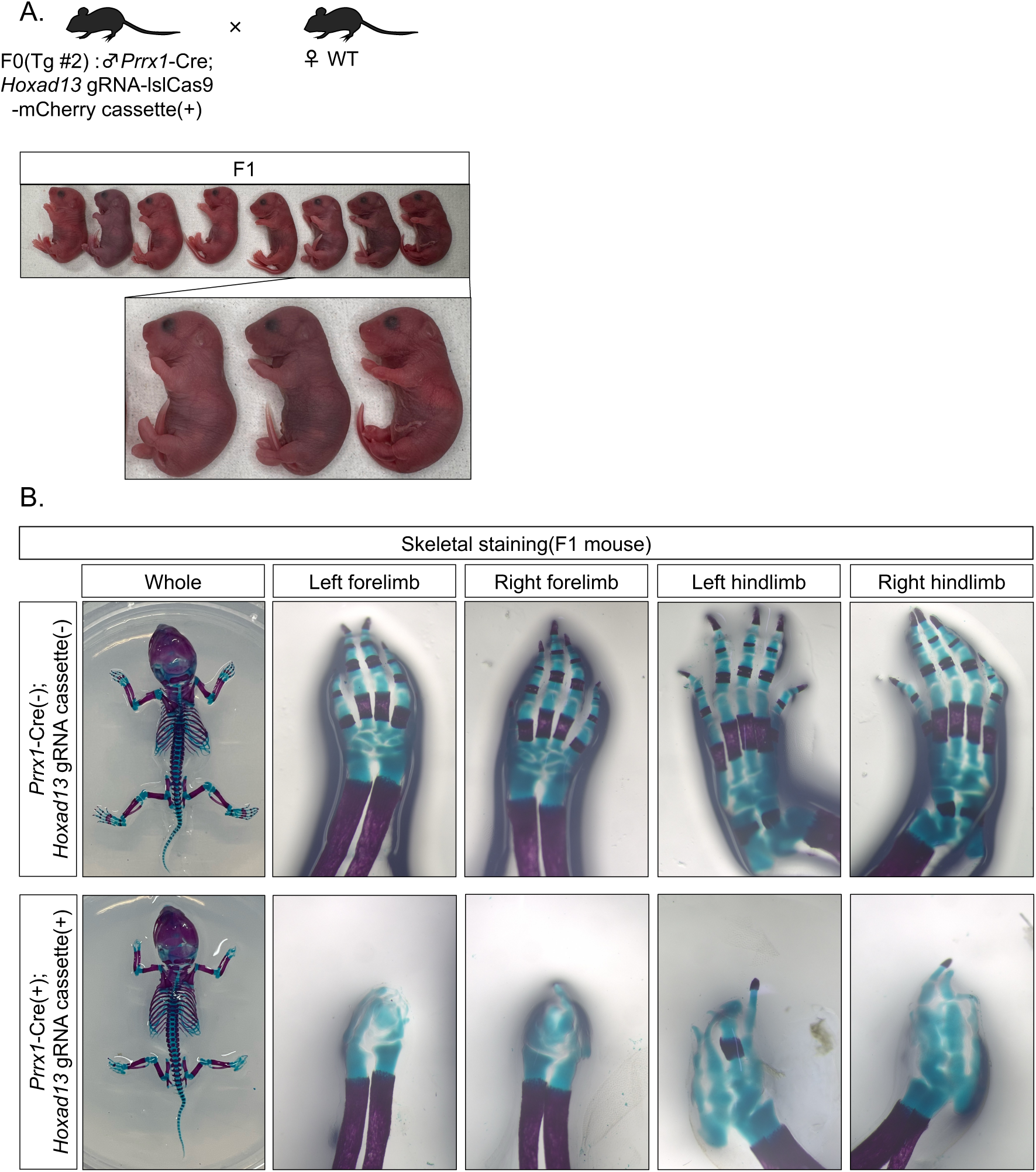
Germline transmission of the conditional *Hoxa13/Hoxd13* gene-edited allele and phenotypic analysis of F1 mice. **(A)** Breeding scheme used for the analysis of germline transmission. An F0 male mouse (Tg#2) carrying the *Prrx1*-Cre transgene and the Hoxa13/d13 gRNA-lslCas9-mCherry cassette was crossed with a wild-type female. Representative F1 offspring are shown. A subset of F1 mice exhibited limb abnormalities similar to those observed in the F0 mouse. **(B)** Skeletal analysis of representative F1 mice. Whole-mount skeletal preparations and enlarged views of the forelimbs and hindlimbs are shown. F1 mice carrying both the *Prrx1*-Cre transgene and the Hoxa13/d13 gRNA cassette exhibited severe autopod malformations. In contrast, control mice lacking the Hoxa13/d13 gRNA cassette displayed normal limb skeletal morphology. Cartilage is stained blue, and bone is stained red.

Skeletal preparations were generated from F1 mice lacking both the *Prrx1*-Cre transgene and the gRNA cassette, as well as from F1 mice positive for both the *Prrx1*-Cre transgene and the gRNA cassette that exhibited incomplete autopod formation. In the forelimb, the autopod elements (carpals, metacarpals, and phalanges) were not observed. In the hindlimb, although some autopod elements (tarsals, metatarsals, and/or phalanges) appeared to be present, they were too poorly defined to be identified with certainty (Fig. 3B).

## Method

### Animals

All mice were maintained under appropriate environmental conditions. C57BL/6JmsSlc mice were purchased from Sankyo Labo Service Corporation, Inc. (Tokyo, Japan). *Prrx1*-Cre mice were obtained from Jackson Laboratory (Bar Harbor, Maine). Pseudopregnant mice (Jcl; ICR, 9–11 weeks old) were purchased from CLEA Japan, Inc. (Tokyo, Japan). All animal experiments were approved by the Institutional Animal Care and Use Committee of Institute of SCIENCE TOKYO.

### Plasmid construction

gRNA cassette vector: A single gRNA expression unit was constructed by placing a guide RNA (gRNA) downstream of the human U6 promoter. Six of these units were then tandemly assembled. Downstream of the gRNA cassette array, a CAG-LSL-Cas9-P2A cassette identical to that used by Feng Zhang and colleagues was inserted. An mCherry fluorescent reporter gene was fused downstream of the P2A sequence. The entire construct was flanked by the 5′ ITR and 3′ ITR piggyBac recognition sequences. IVT donor vector: To enhance RNA stability, the hyPBase coding sequence was flanked by the human β-globin 5′ UTR and 3′ UTR. In addition, a poly(A)100 [pA100] sequence was incorporated. A T7 promoter sequence specified for use with the mMESSAGE mMACHINE™ T7 mRNA Kit with CleanCap™ Reagent AG was introduced upstream of the construct.

### In vitro transcription

The pCFN-T7-AG 5’HBB-FLAG-hyPBase-3’HBB pA100 vector was linearized by digestion with Esp3I (New England Biolabs) at 37℃ for 6 h. The digested product was separated on a 1% agarose gel, and the target band was excised and purified to serve as the DNA template. Purification was performed using the Monarch® DNA Gel Extraction Kit (T1020; New England Biolabs, Ipswich, MA, USA). mRNA was synthesized from the DNA template using the mMESSAGE mMACHINE™ T7 mRNA Kit with CleanCap™ Reagent AG (A57620; Thermo Fisher Scientific, Waltham, MA, USA). The synthesized mRNA was purified using the MEGAclear™ Transcription Clean-Up Kit (AM1908; Thermo Fisher Scientific).

### In vitro fertilization

Male mice used for sperm collection were *Prrx1*-Cre mice aged ≥8 weeks. Sperm were collected from the cauda epididymis of sacrificed mice and incubated in CARD FERTIUP mouse sperm preincubation medium (KYUDO Co., Ltd., Saga, Japan) for 1 h until insemination in a 5% CO₂ incubator at 37℃. For oocyte collection, female C57BL/6JmsSlc mice (3–4 weeks old) were injected with 7.5 U pregnant mare serum gonadotropin (PMSG) (Asuka Animal Health Co., Ltd., Tokyo, Japan) 3 days prior to IVF. After 48 h, 7.5 U human chorionic gonadotropin (hCG) (Asuka Pharmaceutical Co., Ltd., Tokyo, Japan) was administered. Approximately 14 h after hCG injection, oocytes were collected and cultured in CARD MEDIUM® (KYUDO). Three hours after insemination, embryos were washed with mHTF (KYUDO) and cultured until use for microinjection.

Excess oocytes were cryopreserved in 1 M DMSO and DAP213 (KYUDO), thawed, and subsequently used for microinjection in later trials. All procedures were performed in accordance with the mouse assisted reproductive technology manual of Kumamoto University. F1 mice were generated by performing IVF using sperm from F0 male mice. Female C57BL/6JmsSlc mice aged 3–4 weeks were used as oocyte donors. The resulting two-cell embryos were transferred into the oviducts of pseudopregnant recipient females, and pups were obtained 20 days later.

### Micro injection

Microinjection was initiated approximately 6 h after fertilization. An Olympus microscope and a FemtoJet 4i (Eppendorf, Hamburg, Germany) were used for injection. A holding needle (B100-75-10, Sutter Instrument) and an injection needle (World Precision Instruments, Sarasota, FL, USA) were used. For sample preparation, PiggyBac mRNA (6.25 ng/µL) and gRNA-containing vector (30 ng/µL) were diluted in Tris-EDTA buffer (pH 8.0) (06890-54, Nacalai Tesque, Inc., Kyoto, Japan). The mixture was centrifuged at 15,000 rpm at −2℃ for 15 min, and the supernatant was used as the injection solution. Fertilized eggs were transferred to M2 medium (M7167, Merck, MA, USA), and cytoplasmic microinjection was performed into fertilized eggs at the pronuclear stage. Embryos that developed to the 2-cell stage were transferred into the oviducts of pseudopregnant mice, and pups were obtained 20 days later.

### Genotyping PCR

Genomic DNA was extracted from mouse tail tissue. Lysis buffer (50mM Tris HCl (pH8.0), 100mM NaCl, 10mM EDTA, 0.1%SDS) supplemented with Proteinase K recombinant solution (161-28701, FUJIFILM Wako Pure Chemical Corporation, Osaka, Japan) at a final concentration of 100 µg/mL was used, and samples were incubated at 55℃ overnight. An equal volume of Phenol/Chloroform/Isoamyl alcohol (25:24:1) (311-90151, NIPPON GENE Co., Ltd., Tokyo, Japan) was added to extract DNA, followed by purification via isopropanol precipitation (166-04831, FUJIFILM Wako). PCR was performed using Quick Taq® HS DyeMix (DTM-101, TOYOBO, Osaka, Japan). The PCR products were analyzed by agarose gel electrophoresis. The primers used are listed below.

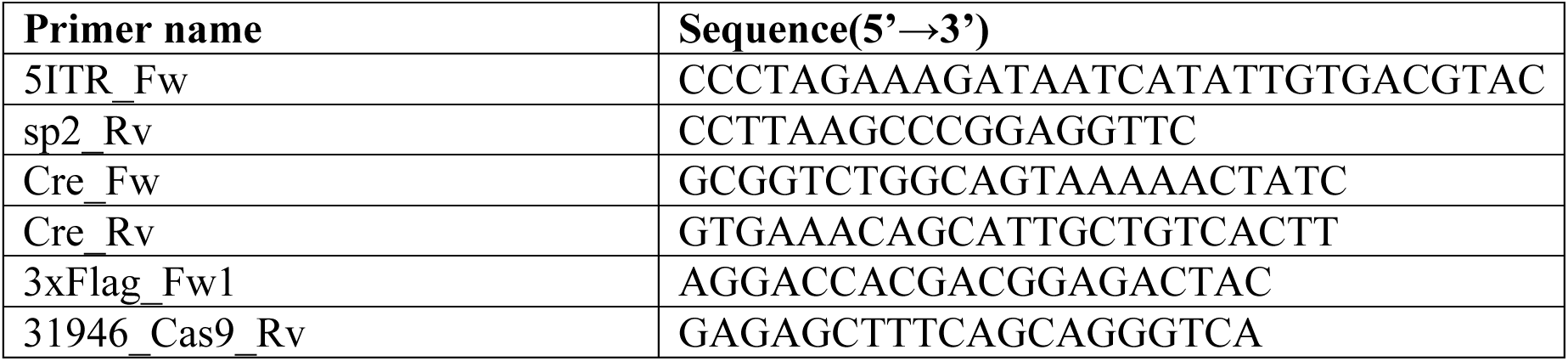

### Skeletal staining specimens

Skeletal staining was performed according to the method reported by Suzuki et al. (2025) [35]. P0 mice were euthanized, frozen, and subsequently thawed in water at 40°C. After thawing, the epidermis was carefully removed under water. A small portion of the epidermis was retained for genotyping analysis. The specimens were fixed and dehydrated in 90% ethanol for 15 min. Adipose tissue, skin, and internal organs were removed in 90% ethanol. The samples were then incubated in 99.5% ethanol at 4°C overnight.

Subsequently, the mice were immersed in acetone for 2 h, followed by replacement with fresh acetone and incubation overnight on a gentle shaker. The specimens were stained in 0.015% Alcian Blue solution (20% acetic acid, 80% ethanol) for two consecutive overnight incubations. After staining, samples were washed in 95% ethanol for 1 h and further dehydrated in 99.5% ethanol with gentle shaking. Samples were then stained in 0.0125% Alizarin Red solution prepared in 70% ethanol for 2 h on a shaker. The specimens were transferred to 1% KOH and incubated at 4°C for 1 h. The solution was replaced with fresh 1% KOH, and samples were further incubated overnight at 4°C. Thereafter, specimens were cleared by sequential incubation in 0.8% KOH/20% glycerol, 0.6% KOH/40% glycerol, 0.4% KOH/60% glycerol, and 0.2% KOH/80% glycerol, with solution changes every other day, followed by storage in 100% glycerol. Images were captured using a stereomicroscope (SMZ800N, Nikon, Tokyo, Japan) equipped with a digital camera (4K-LITE FL, RELYON, Tokyo, Japan).

## Discussion

We combined the transposon, CRISPR/Cas9, and Cre/loxP systems and tandemly linked multiple gRNAs, enabling the generation of multi-gene-modified mice through a single genome-editing event. In this study, we used TEMPO-editing to generate double-gene-edited mice for *Hoxa13* and *Hoxd13*, two genes involved in limb development. A vector carrying lsl-Cas9-mCherry, together with three gRNAs targeting *Hoxa13* and three gRNAs targeting *Hoxd13*, was constructed and introduced into *Prrx1*-Cre fertilized eggs, which express Cre recombinase specifically in the limbs, along with piggyBac mRNA. As a result, the *Hoxa13*/*Hoxd13*_gRNA-lslCas9-mCherry cassette was excised from the vector by the piggyBac transposase and successfully integrated into the genome. In the presence of Cre recombinase, Cas9 was subsequently expressed, resulting in genome editing of the target genes. Consistent with previous reports [22], this approach successfully recapitulated the absence of autopod formation. Among the F0 mice obtained, insertion of the gRNA cassette(s) was confirmed in 9–25% of the offspring.

The TEMPO-Editing approach developed in this study represents an improvement of our previously reported one-step method for generating genetically modified mice [13] in two major aspects. First, whereas the conventional method employed only a single gRNA cassette for each target gene, TEMPO-Editing was designed to simultaneously express three distinct gRNAs per gene. This strategy enables the induction of large genomic deletions within the target locus, thereby increasing the efficiency of gene disruption and resulting in more robust gene-edited mice generation. Second, in the conventional method, a gRNA-expressing cassette vector was injected into fertilized eggs obtained from crosses between Cre-transgenic mice and lsl-Cas9 knock-in (KI) mice [9]. Therefore, two separate mouse lines—the Cre line and the lsl-Cas9 KI line—had to be maintained and prepared for experiments. In contrast, TEMPO-Editing requires only a single mouse line, namely a male mouse carrying the desired Cre transgene, greatly simplifying the experimental procedure and substantially reducing the effort required for the generation of genetically modified mice. Although six gRNAs were tandemly linked in the present study, TEMPO-Editing has the potential for much greater multiplexing. In fact, we have confirmed that nearly 40 gRNAs can be inserted into the genome as a single cassette (unpublished data).

Furthermore, the *Hoxa13*/*Hoxd13* double-gene-edited mice generated in this study could be used to investigate the roles of these genes in adulthood by simply changing the Cre driver line. Increased *HOXA13* expression has been reported in esophageal adenocarcinoma [24], and mutations in *HOXA13* are known to cause developmental defects of the female reproductive tract [25]. In addition, *HOXA13* has been suggested to function within a developmental pathway shared with Wnt5 during uterine morphogenesis [26]. In contrast, aberrant *HOXD13* expression has been reported in human hematological malignancies [27]. Moreover, Hox genes that regulate skeletal patterning during embryonic development are known to be reactivated following bone fracture or tissue injury in adulthood [28]. To date, however, no studies have reported the use of *Hoxa13*/*Hoxd13* double conditional gene-edited mice to investigate the functions of these genes at the adult stage. Therefore, determining the physiological consequences of simultaneous *Hoxa13* and *Hoxd13* deficiency in adult animals represents an important research objective that may reveal previously unrecognized functions of Hox genes. In addition, the adult functions of the Hox13 paralog group, consisting of *Hoxa13*, *Hoxb13*, *Hoxc13*, and *Hoxd13*, remain poorly understood. We anticipate that TEMPO-Editing will enable the generation of Hox13 quadruple-gene-edited mice, providing a powerful strategy for elucidating the collective functions of these genes in adult physiology. Thus, this system enables the generation gene-edited mice targeting entire gene families or all candidate genes identified through screening within a remarkably short period of approximately one month. Although a major advantage of this system is that phenotypic analyses can be performed directly in the F0 generation, we have also confirmed that the inserted cassettes are efficiently transmitted to subsequent generations, making it possible to establish and maintain stable mouse lines.

In addition, we believe that the gRNA-containing cassettes developed in this study can, in principle, be applied to any experimental animal species in which piggyBac transposon-mediated genome engineering is feasible. Notably, successful piggyBac-based genome modification has been reported in several model organisms, including non-human primates such as cynomolgus monkeys, zebrafish, and insect models [29–31]. Therefore, TEMPO-Editing has the potential to serve as a versatile platform for rapid and efficient generation of multiplex genome-edited animals across a broad range of species.

Genome editing technologies that combine transposon systems with CRISPR/Cas9 have recently been applied beyond conventional gene knockout approaches. Notably, Hatada and colleagues reported that this strategy enables the highly efficient generation of epigenome-edited mice, which have traditionally been difficult to establish [32]. Epigenetic abnormalities are known to contribute to a variety of hereditary diseases, and conventional approaches often suffer from low transgenic production efficiency. Moreover, when epigenetic modifications induce severe phenotypes, animals may die or become infertile before F1 offspring can be obtained. To overcome these limitations, Hatada et al. developed a system in which a large cassette containing dCas9, epigenetic modifier(s), a gRNA expression cassette, and a reporter gene is introduced using a transposon-based strategy. This approach enables direct phenotypic and molecular analyses in the F0 generation, thereby bypassing the need for extensive breeding and substantially accelerating the generation and evaluation of epigenome-edited animal models.

In recent years, genome-editing technologies based on CRISPR-associated nucleases have rapidly advanced. In particular, next-generation CRISPR systems such as Cas12, which is well suited for multiplex genome editing and exhibits lower off-target activity [33], and Cas3, which is capable of inducing large deletions spanning tens of kilobases [34], have emerged as promising alternatives to conventional Cas9-based approaches. We are also interested in investigating whether combining genome editing approaches based on these next-generation Cas enzymes with transposon-mediated editing technologies could enable the development of even more efficient and versatile genome engineering platforms.

Overall, the TEMO-Editing system developed in this study represents a powerful tool for the simpler and more rapid generation of conditional gene-edited mice. We anticipate that this method will lower the barriers to in vivo functional analyses across a wide range of research fields and thereby contribute to further advances in biological research.

## Acknowledgments

We are grateful to all staff of the Department of Systems BioMedicine at Institute of SCIENCE TOKYO(IST) for their support. We are grateful for the help and support provided by the Animal Research Facilities, Bioscience Center, Research Infrastructure Management Center. This work was supported by Japan Agency for Medical Research and Development (AMED) (Grant Numbers JP22gm0010009, JP22ama121045, JP24jf0126010, JP24gm2010002, JP25ek0109836), Japan Society for the Promotion of Science (JSPS) KAKENHI (Grant Numbers JP20H05696), JST Program for co-creating startup ecosystem(Grant Number JPMJSF2313), Japan, to H.A.

## Author Contributions

M.I. and Y.K. performed in vitro fertilization (IVF), microinjection, embryo transfer, and genomic analysis, and wrote the manuscript. S. M. selected gRNAs and constructed the plasmid. T. C. made the concept of this study, selected gRNAs and constructed the plasmid. M. T. performed genomic analysis. H.A. supervised this project.

## Notes

### Competing Interest Statement

The authors have declared no competing interest.

